# Efficacy of auxin-inducible protein degradation in *C. elegans* tissues using different auxins and TIR1-expressing strains

**DOI:** 10.1101/2024.01.16.575916

**Authors:** Nidhi Sharma, Filipe Marques, Paschalis Kratsios

**Affiliations:** Department of Neurobiology, University of Chicago, Chicago, IL, USA

## Abstract

The auxin-inducible degradation system has emerged as a powerful tool to deplete proteins of interest in cells and tissues of various model organisms, including *C. elegans* ^2–5^. Here, we present a detailed protocol to perform AID-driven spatiotemporal depletion of specific proteins in *C. elegans* tissues. First, we introduced the AID degron and a fluorescent reporter at two conserved proteins: (a) the transcription factor CFI-1 (human ARID3), which is expressed in the nucleus of multiple *C. elegans* neurons and head muscle cells ^6,7^, and (b) the broadly expressed translation initiation factor Y47D3A.21 (human DENR) that localizes in the cytoplasm. Second, we provide a step-by-step guide on how to generate *C. elegans* strains suitable for AID-mediated protein (CFI-1 and DENR) depletion. Third, we find that the degree of CFI-1 and DENR depletion in *C. elegans* tissues is comparable upon treatment with either natural auxin (indole-3-acetic acid (IAA) or a water-soluble synthetic auxin analog (K-NAA). Last, we compare the degree of AID-mediated CFI-1 depletion in *C. elegans* neurons when the transport inhibitor response 1 (TIR1), component of the SCF ubiquitin ligase complex, is provided in neurons or all somatic cells. Altogether, this protocol provides side-by-side comparisons of different auxins and TIR1-expressing lines. Such comparisons may benefit future studies of AID-mediated protein depletion in *C. elegans*.

**Graphical abstract:** **Image provided as pdf (together with Figures).**

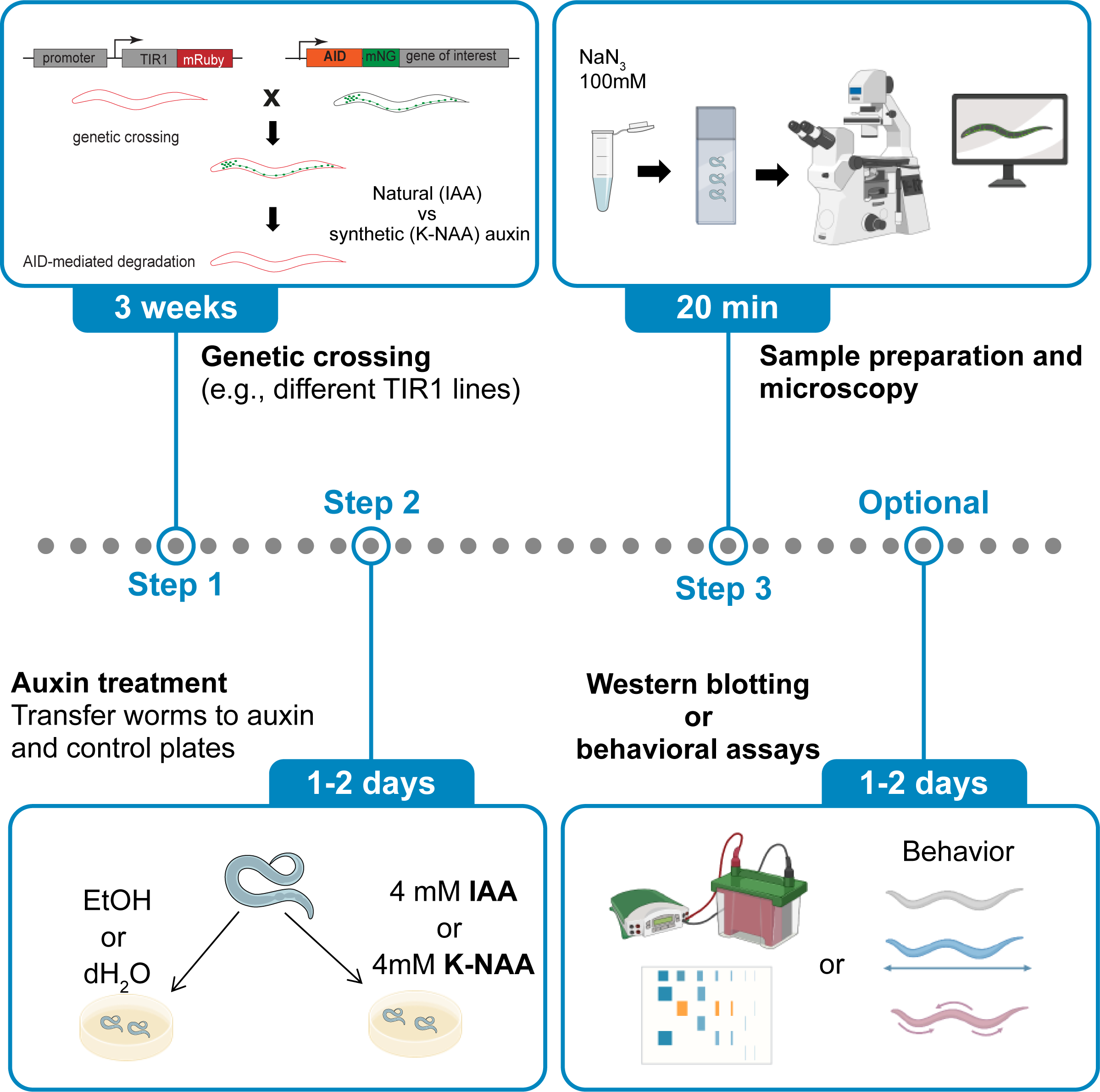

**Highlights:**

- Efficient protein depletion in *C. elegans* tissues upon treatment with either natural or synthetic auxins.
- Pansomatic TIR1 expression leads to efficient depletion of CFI-1 and DENR.
- Panneuronal TIR1 expression leads to neuron-specific, yet variable CFI-1 depletion.
- The AID system is compatible with fluorescence microscopy, Western blotting and behavioral assays.

## Before you begin

We detail in this protocol the steps necessary for achieving auxin-inducible protein depletion in *C. elegans* tissues. As proof-of-principle, we focus on two conserved proteins with different subcellular localization: (a) the transcription factor CFI-1 (human ARID3) is expressed in the nucleus of *C. elegans* neurons and head muscle cells^6–8^, and (b) the ubiquitously expressed translation initiation factor Y47D3A.21 (human DENR) that localizes in the cytoplasm. The AID system is tripartite and requires (1) the plant hormone auxin or a synthetic analog, the protein of interest must be tagged with the auxin inducible degron, and (c) the *Arabidopsis thaliana* F-box protein, transport inhibitor response 1 (TIR1), must be expressed in the cell type or tissue of interest (**Figure 1A**)^4^. Upon exposure to auxin, TIR1 acts as a substrate to ubiquitinate the AID-tagged protein, enabling its proteasomal degradation. The AID sequence can be introduced into the endogenous locus of the gene of interest by CRISPR/Cas9 gene editing, as detailed in a recent protocol ^2^. With this method, we successfully tagged CFI-1 with a green fluorescent protein (mNeonGreen) fused to AID and Y47D3A.21 (DENR) with GFP::AID (**Figure 2**). Next, we obtained two transgenic strains that express TIR1 either in all *C. elegans* somatic cells or exclusively in neurons (see Key Resources Table) (**Figure 1B**). Through genetic crossing and/or CRISPR/Cas9 gene editing, we generated homozygous animals for the AID allele (e.g., *cfi-1(kas16[mNG::AID::cfi-1]) I)* and the TIR1 transgenes. The steps we detail here for CFI-1 and Y47D3A.21 can be applied to any *C. elegans* protein.

**Figure 1:**
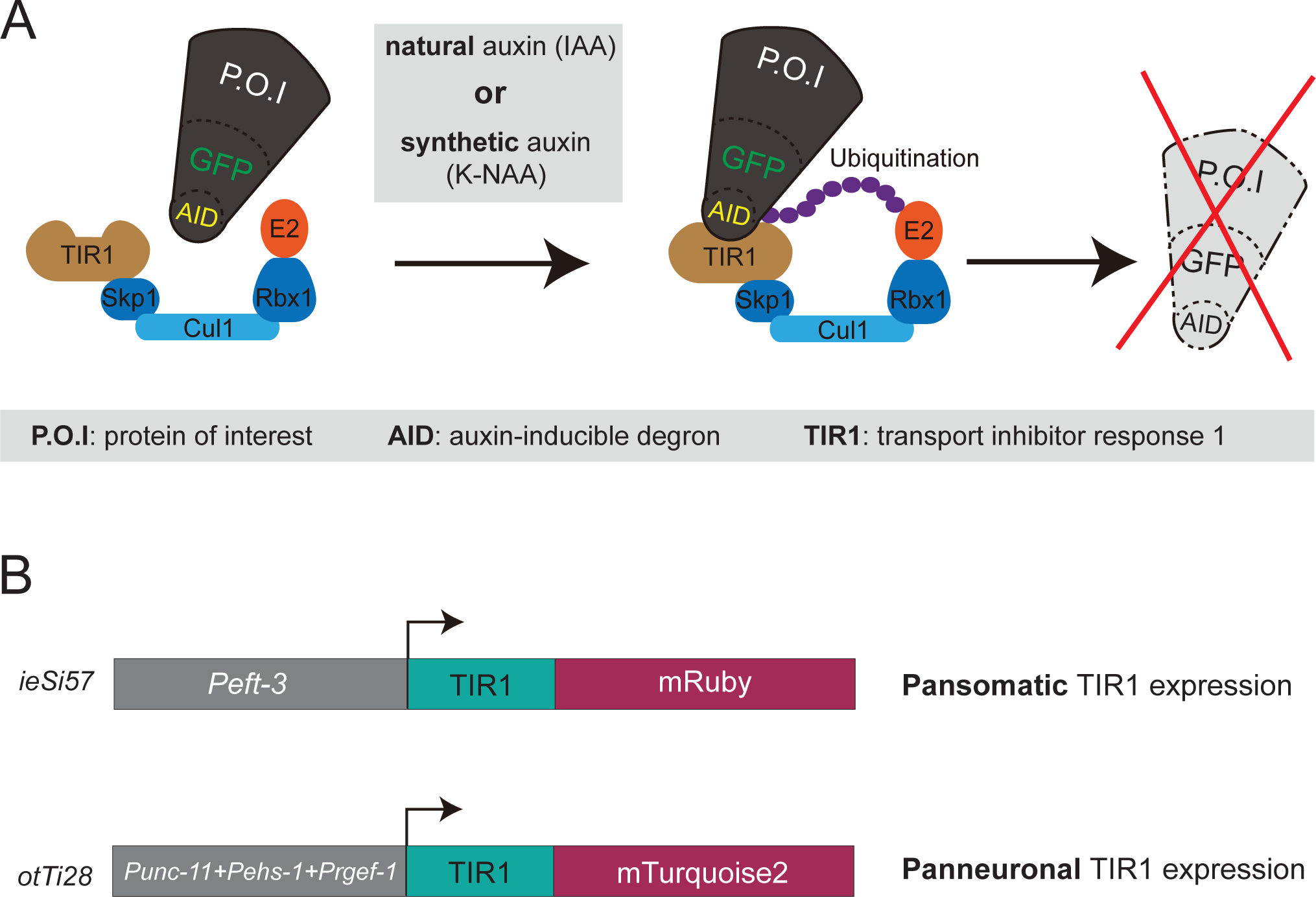
AID-mediated protein depletion using different auxins and TIR1-expressing lines. **A:** Schematic of auxin-inducible degradation system. Natural (IAA) or synthetic (K-NAA) auxin treatment was employed in this protocol. **B:** Schematic of the two TIR1 lines used in this protocol.

**Figure 2:**
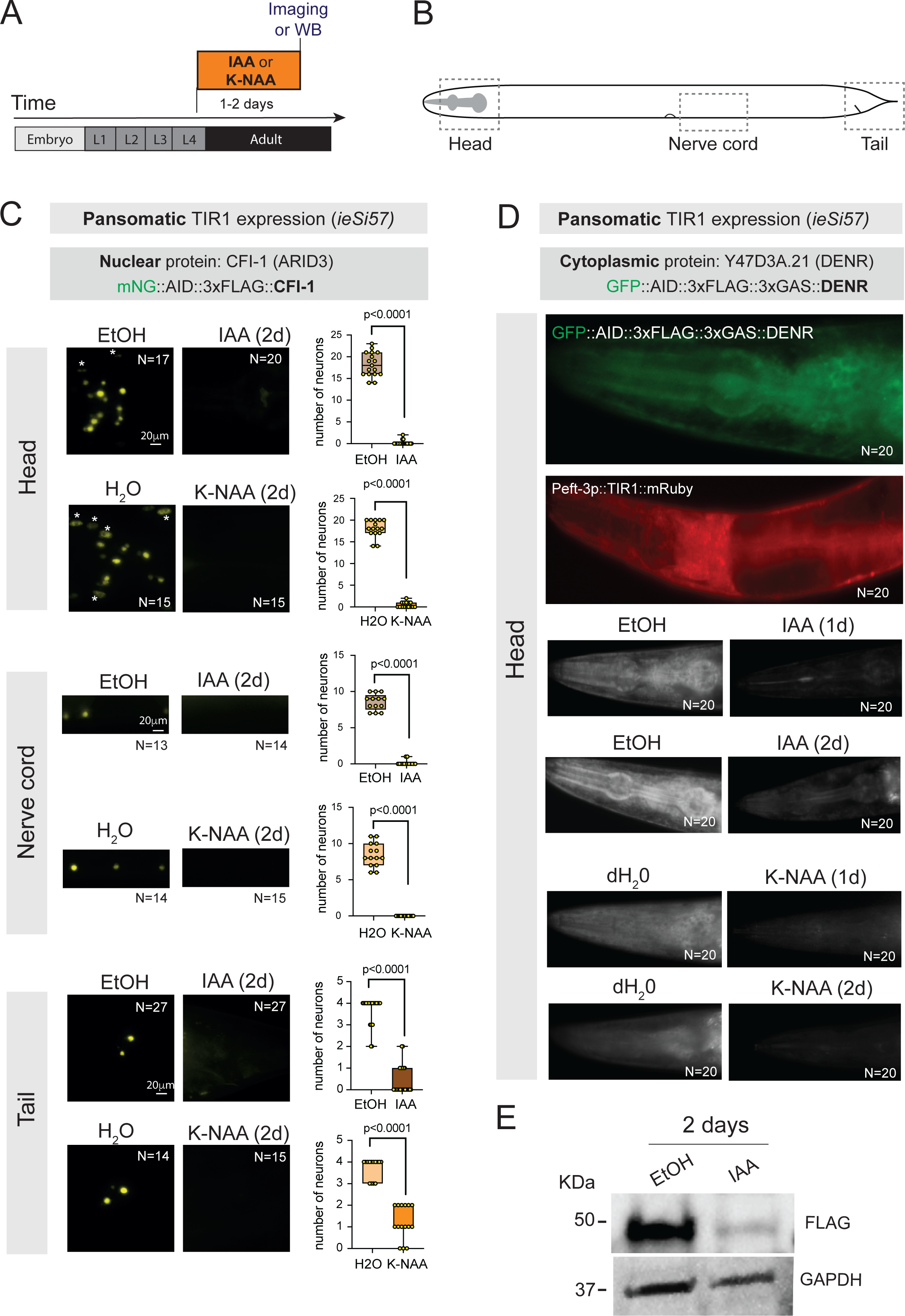
AID-mediated depletion of nuclear and cytoplasmic proteins using two different auxins. **A:** Schematic of auxin treatment. **B:** Schematic of *C. elegans* body. Boxed regions indicate images shown in panels C and D. **C**: Representative images showing mNG::AID::3xFLAG::CFI-1 expression in head neurons and head muscle cells (asterisks) upon IAA or K-NAA treatment for two days. Controls: Two day-long exposure to either 100% ethanol or dH2O. Strain genotype: *cfi-1(kas16[mNG::AID::cfi-1]) I; ieSi57 [Peft-3::TIR1::mRuby::unc-54 3’ UTR, cb-unc-119(+)] II*. Images from the *C. elegans* head, ventral nerve cord, and tail are shown. Quantification of the number of neurons expressing mNG is provided on the right. Student’s t test, N.S: Not significant. N: number of animals. **D**: Representative images showing *<u>GFP-AID-3Xflag-3xGAS::</u> Y47D3A.21 and Peft-3::TIR1::mRuby expression in the C. elegans head region (top panels).* Expression of <u>GFP::AID::3XFLAG::3xGAS::</u> Y47D3A.21 in the cytoplasm of most cells and tissues is observed. Representative images are shown upon IAA or K-NAA treatment for two days. Controls: Two day-long exposure to either 100% ethanol or dH2O. **E**: Western blotting with antibodies against FLAG and GAPDH on homozygous <u>GFP::AID::3XFLAG::3xGAS::</u> Y47D3A.21 animals with pansomatic TIR expression upon IAA treatment for two days.

Since its initial establishment in *C. elegans* in 2015 ^4^, the AID system has continued to evolve ^2,9–11^. A battery of cell type-specific TIR1-expressing strains is now available, enabling auxin inducible protein depletion in specific *C. elegans* tissues (e.g., germline, intestine, nervous system, muscle) ^12^. Because natural auxin (indole-3-acetic acid [IAA]) has several disadvantages (light-sensitivity, no penetrance of eggshell, requires ethanol as a solvent), synthetic auxin analogs, such as 1-naphthaleneacetic acid, potassium salt (K-NAA) and 5-phenyl-indole-3-acetic acid (5-Ph-IAA), have been developed ^2,3,11,13,14^. However, a side-by-side comparison of depleting the same protein using different TIR1 lines, as well as natural and synthetic auxins, is lacking in *C. elegans*.

In this protocol, we first outline all necessary steps for achieving AID-mediated protein depletion in *C. elegans*, ranging from strain generation to auxin administration. Further, we compare the efficacy of natural auxin (indole-3-acetic acid (IAA) and a water-soluble synthetic auxin analog (K-NAA) in depleting a nuclear (CFI-1) and a cytoplasmic (Y47D3A.21[DENR]) protein (**Figure 1A**). Last, we directly compare the degree of AID-mediated CFI-1 depletion in *C. elegans* tissues when TIR1 is provided only in neurons or all somatic cells(**Figure 1B**).

### Genetic crossing of *C. elegans* strains

#### Timing: ∼3 weeks

1. Genetic crossing entails the following steps:

a. Male animals for the AID allele (e.g., *cfi-1(kas16[mNG::AID::cfi-1]) I)* can be generated by crossing into N2 control hermaphrodites or via heat-shock exposure.
b. Fifteen malesthat carry the AID allele are crossed with 5 hermaphrodite animals homozygous for the TIR1-expressing transgene.
c. At the next generation, pick 5 hermaphrodites that are heterozygous for both the AID allele and TIR1 transgene.
d. Allow worms to generate self-progeny and continue to pick and single 5 hermaphrodites until both the AID allele and the TIR1 transgene become homozygous.

**CRITICAL**: It is important to use only well-fed (not starved animals) for genetic crosses. Set up crosses by picking animals at larval stage 4 (L4).

Note: As described above, we introduced the TIR1 transgene via genetic crossing to *cfi-1(kas16[mNG::AID::cfi-1])* animals. However, for the cytoplasmic protein (DENR), we employed CRISPR/Cas9 gene editing to generate the *Y47D3A.21(syb6129[<u>GFP-AID-3Xflag-</u> <u>3xGAS::</u> Y47D3A.21])II allele following procedures described in* ^2^*. Gene editing was performed on homozygous animals for the pansomatic TIR1 line (ieSi57)*.

### Preparation of Solutions

#### Timing: 2 h

2. Fresh preparation of nematode growth medium (NGM). The NGM composition is:

a. NaCl, Agar, Peptone, 5mg/ml Cholesterol (prepared in 100 % ethanol), 1M MgSO_4_, 1M KPO_4_ buffer pH 6.0.
b. 10mm petri dish.
3. Fresh preparation of solutions containing natural auxin (indole-3-acetic acid (IAA):

a. The natural auxin indole-3-acetic acid (IAA) was purchased and stored in –20 °C up to a year or recommended by manufacturer otherwise.
b. 100 % ethanol is used as IAA dissolvent.
c. A 400 mM stock solution in ethanol was prepared and stored at 4 °C for up to one month.
4. Fresh preparation of solutions containing a water-soluble synthetic auxin analog (K-NAA):

a. The water-soluble synthetic auxin analog (K-NAA) was purchased and stored in –20 °C up to a year or recommended by manufacturer otherwise.
b. Deionized water (dH_2_O) is used as auxin dissolvent.
c. A 400 mM stock solution in dH2O was prepared and stored at 4 °C for up to one month.

**CRITICAL**: The auxin (IAA) solution should be protected from light exposure and shielded with aluminum foil. However, a recent protocol indicates that the photostability of synthetic auxin (K-NAA) prevents light-induced compound degradation during storage ^3^.

**Note**: Check reagents details in the Key Resources Table.

### Preparation of Auxin-NGM plates and seeding with food source (*E*. *Coli* OP50 strain)

#### Timing: 2-3 Days

5. NGM plate preparation (∼ 1 Day) (Supplementary Figure 1).

a. Autoclaved NGM was cooled down to 50°C and split into two bottles of equal volume (500 ml) for Auxin and ethanol (negative control).
b. Add 5 ml of 400 mM auxin (IAA or K-NAA) to 500 ml of NGM to the final concentration of 4 mM.
c. As a control for natural auxin (IAA), add 5 ml of 100% ethanol into 500 ml of NGM.
d. As a control for synthetic auxin (K-NAA), add 5 ml of dH2O into 500 ml of NGM.
e. Pour 10ml of prepared auxin-NGM and control-NGM to 55mm petri dish and let it dry for one day.
6. Seeding OP50 bacteria on auxin and ethanol coated NGM plates (∼ 2 Days).

a. add 200 µl of concentrated OP50 on NGM plates and let it dry for 2 days.

**CRITICAL**: Before adding auxin, the temperature of NGM should be 50°C. Otherwise, it may cause a rapid degradation of auxin. In addition, auxin (IAA, K-NAA) concentration in the NGM plate may vary from 500 mm to 4 mM; 4 mM is the highest concentration used in this protocol. However, we recommend optimization of auxin concentrations for each experiment.

**Note**: NGM plates containing auxin (IAA or K-NAA) can be stored at 4 °C up to two weeks. In our experiments, we shielded both NGM IAA and NGM K-NAA plates from light (Supplementary Figure 1). However, a recent protocol indicates that the photostability of K-NAA prevents light-induced compound degradation during storage ^3^.

## Key Resources Table

| REAGENT or RESOURCE | SOURCE | IDENTIFIER |
| --- | --- | --- |
| Bacterial and virus strains |  |  |
| OP50 Escherichia coli | Caenorhabditis Genetics Center (CGC) | OP50 |
| Chemicals, peptides, and recombinant proteins |  |  |
| Agar, Bacteriological | Lab Scientific bioKEMIX | A466 |
| NaCl | SIGMA | 71376-1KG |
| Peptone | Thermo Fisher Scientific | BP1420-2 |
| CaCl<sub>2</sub> | ACROS | CAS 10043-52-4 |
| MgSO<sub>4</sub> | Sigma Aldrich | M7506-500G |
| Potassium phosphate, monobasic | ACROS | CAS 7778-77-0 |
| Cholesterol | Sigma Aldrich | C8667-5G |
| Indole-3-acetic acid, 98+% (Auxin) | Alfa-Aesar | CAS 87-51-4 |
| 1-naphthaleneacetic acid, potassium salt (K-NAA) |  |  |
| Tryptone | Sigma Aldrich | T-7293-250G |
| Yeast Extract | SIGMA | Y1625-250G |
| Sodium Azide | ACROS | CAS 26628228 |
| Agarose (Protein Electrophoresis Grade), Fisher BioReagents™ | Fisher Scientific | BP164-500 |
| Ethanol 200 Proof | DECON LABS | 2701 |
| Experimental models: Organisms/strains |  |  |
| *C. elegans*: *cfi-1(kas16[mNG::AID::cfi-1]) I* | This study | KRA345 |
| *C. elegans*: *otTi28 [unc-11prom8+ehs-1prom7+rgef-1prom2::TIR1::mTurquoise2::unc-54 3'UTR] X* | Hobert lab (Columbia University) | OH14930 |
| *C. elegans*: *unc-119(ed3); ieSi57 [Peft-3::TIR1::mRuby::unc-54 3' UTR, cb-unc-119(+)] II* | Caenorhabditis Genetics Center (CGC) | CA1200 |
| *cfi-1(kas16[mNG::AID::cfi-1]) I; ieSi57 [Peft-3::TIR1::mRuby::unc-54 3' UTR, cb-unc-119(+)] II* | This study | KRA830 |
| *cfi-1(kas16[mNG::AID::cfi-1]) I; otTi28 [unc-11prom8+ehs-1prom7+rgef-1prom2::TIR1::mTurquoise2::unc-54 3'UTR] X* | This study | KRA842 |
| *C. elegans*: *Y47D3A.21(syb6129[GFP-AID-3Xflag-3xGAS::Y47D3A.21])III; ieSi57II* | This study | PHX6129 |
| Software and algorithms |  |  |
| Version 2.3.69.1000, Blue edition | Carl Zeiss Microscopy | https://www.zeiss.com/microscopy/en/products/software/zeiss-zen.html |
| Other |  |  |
| Dissection microscope | Carl Zeiss | Stemi 305 |
| Glass Microscopic Slides (Plain) | Fisher Scientific | 12-544-1 |
| Falcon™ Bacteriological Petri Dishes with Lid (55mm) | Fisher Scientific | 08-757-100B |
| Zeiss, Axio Imager Z2 | Carl Zeiss Microscopy | https://www.zeiss.com/microscopy/en/products/light-microscopes/widefield-microscopes/axio-imager-2-for-life- |
|  |  | science-research.html |
| Aluminum Foil |  |  |

## Materials and equipment

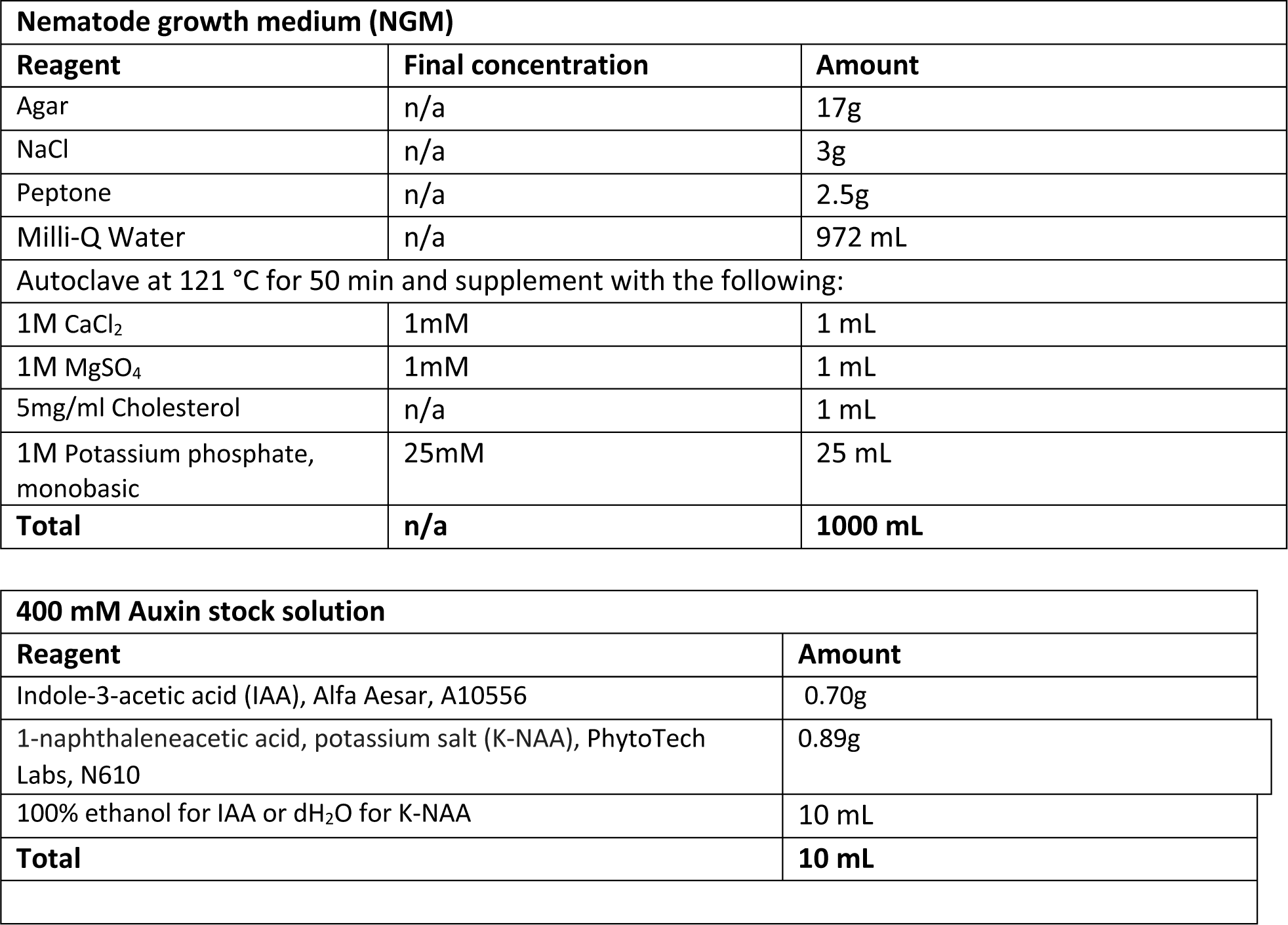

[Storage condition: at 4 °C and shielded from light for up to one month although fresh preparation each time is recommended for better results]

## Step-by-step method details

### [Auxin treatment]

#### Timing: [2 days]

This step requires transferring the worms onto NGM plates with ethanol (EtOH) or 4mM natural auxin (IAA), or onto NGM plates with dH_2_O or 4mM synthetic auxin (K-NAA). The duration of the auxin exposure depends on the protein of interest. Keep all NGM IAA plates shielded from light by covering them with aluminum foil.

1. Transfer worms to treatment plates.

a. Pick worms with the platinum wire picker at L4 (or younger). N = 25 or higher.
b. Transfer them to bacteria seeded NGM-auxin (4mM) and NGM-ethanol plates.
2. Incubate the worms for 1-2 days at 20° C. **See problem 1 and 2.**

**CRITICAL:** It is important to shield the NGM IAA plates from light because natural auxin (IAA) is light sensitive. Further, worms grown on auxin plates may present a slight delay in growth. This is particularly important if you need to image worms at a specific developmental stage. Because dH_2_0 (not ethanol) is used to dissolve K-NAA, the worms on K-NAA NGM plates appear healthier compared to worms on IAA (dissolved in 100% ethanol) NGM plates. The solvent of IAA (ethanol) may confound *C. elegans* lifespan and behavioral assays.

### [Sample preparation for fluorescence microscopy]

#### Timing: [15 min]

This step involves the preparation of anesthetized worms for imaging and mounting them on a 4% agarose pad on glass slides. These pads prevent worms from being crushed between the coverslip and glass slide, and help avoid desiccation.

3. Prepare 4% agarose solution.

a. Add 4 g agarose to 100 ml dH2O in a 250-ml Erlenmeyer flask.
b. Heat up, very carefully, in a microwave until all agarose dissolves.
c. Place the melted agarose on a heat block, at approximately 95°C, while preparing the slides, to prevent solidification.

**Optional**: Alternatively, agarose can be dissolved on a stirring hotplate, with a stir bar.

**Note**: The 4% agarose solution can be reused. Microwave upon solidification.

**Pause point**: Turn on microscope (e.g., Zeiss Axio Imager Z2), computer, laser source and associated equipment before starting to prepare the samples.

4. Prepare the imaging slides (Supplementary Figure 2).

a. Prepare two support slides by placing two layers of laboratory tape on each slide. Place these support slides on each side of a blank side.
b. Place one drop (∼600 μL) of warm 4% agarose onto the blank slide using a laboratory pipette with a cut blue tip.
c. Quickly place a second blank slide on the top of the agarose drop, at a 90° angle to the three slides, and gently press the edges of this slide such that the agarose spreads uniformly. Let the agarose dry for a minute.
d. Separate the glass slides without perturbing the agarose pad. Use a razor blade to trim excessive parts and adjust the agarose pad.
e. The control and treatment samples can be mounted on the same slide by dividing the agar pad into several sections. The agar pad can be divided into 9 smaller squares if done carefully, allowing faster imaging since the slides do not have to be changed out between each condition.

**Note**: The support slides ensure that the pad thickness (∼0.2mm) will remain the same across all samples.

**Optional**: A glass Pasteur pipette can be used to drop the warm agarose on the blank slide.

**Note**: Prepare fresh agarose pads for each imaging session

5. Mount the worms.

a. Place a 10 μL droplet of 100mM sodium azide (NaN_3_) on the agarose pad. **See problem 5.**
b. Pick the worms to be imaged under the dissection scope and put them in the NaN_3_ droplet. **See problem 6.**
c. Use the platinum wire picker to move the worms around within the droplet so that they do not overlap with each other.
d. Gently cover the droplet with a coverslip. The worms are ready for imaging. **See problem 7**.

**Note**: This droplet volume of anesthetizing agent is sufficient for 10 to 20 worms but respectively increase or decrease the volume if more or less worms are analyzed. **See problem 8.**

### [Microscopy and image acquisition]

#### Timing: [15 min]

This step includes the microscope set-up and image acquisition.

Images can be taken with an automated fluorescence microscope (e.g., Zeiss Axio Imager Z2) and a camera (e.g., Zeiss Axiocam 503 mono) using the ZEN software (Version 2.3.69.1000,Blue edition).

6. Turn on the microscope and associated equipment.

a. Turn the power on for the laser source.
b. Turn on the computer.
c. Open software for image acquisition.
7. Once calibration of the microscope system is complete, begin imaging samples. Recalibrate every timing the system is restarted.
8. Place the imaging slide that contains the sample on the microscope stage with the glass coverslip facing the objective lens.
9. Turn the bright-field light source on and direct light to the eyepiece of the microscope.
10. At the lowest magnification, gradually adjust the position of the stage to bring the sample to the center of view, search the worms and then focus using coarse or fine focus adjustment. **See problem 9.**
11. Switch to the desired magnification and re-focus the sample.
12. Switch to the desired laser filter

a. Adjust the exposure time to your fluorescent reporter and avoid saturation.
b. Avoid photobleaching by minimizing the time that a worm is exposed to light.
13. Acquire Z-stack images with 0.5-1 μm intervals between stacks.
14. Save all images and carefully remove the slide.
15. Once all samples have been imaged turn off all the equipment.

**Note**: Allow approximately 20 minutes for the laser to power up and equilibrate before image acquisition.

**Note**: The optimal thickness is ∼0.25 μm (recommended by Zen software), however that will increase the stack size and acquisition time, as well as the image size.

**Note**: The imaging parameters might need to be adjusted depending on the specific platform used. Some microscopes require the user to specify the exact wavelength if using a white light laser, and those using advanced detectors might also have to specify the emission range for detection.

**Note**: Generally, at least 20 worms should be analyzed for each condition.

**Critical**: The same imaging parameters should be used across all samples throughout the experiment. Any change can significantly alter signal strength and complicate results interpretation.

## Quantification analysis of fluorescent microscopy signals

Images can be obtained using ZEN software (Zeiss) or any other open-source alternative such as FIJI/ImageJ^15,16^. This section includes guidance on quantification of fluorescent microscopy signals using FIJI/ImageJ (**Supplementary Figure 3**), an open-source software.

1. Open the image files to be quantified in ImageJ. Go to tab “File → Open” to select files individually, or select all files (in Windows, use “Control” plus mouse click; in Mac OS, use “Command” plus mouse click), drag the files, and drop into the ImageJ window.
2. Choose the parameters to be analyzed under the tab ”Analyze → Set Measurements…”. Check the desired parameters and click OK to close the window.

a. Typical parameters to be measured include “Area”, “Mean gray value”, “Min and Max gray value”, ‘Standard deviation” and “Integrated density”.
3. Use a background subtraction method to enhance the differences between similar images and to remove undesired signaling from the background. Go to tab “Process → Subtract background…”. Check Sliding paraboloid, keep the rolling ball value at 50 pixels and click OK to close the window.
4. If a Z-stack image was acquired go to tab “Image → Stacks → Z project…”. Select Max Intensity as projection type and click OK to close the window. A new image window will show up with your maximum projection.
5. In the ImageJ window, click on the freehand selection tool. On the maximum projection image, carefully outline the region to be quantified.

a. If it helps, the image can be zoomed in to get a closer look when outlining the region of interest. Go to “Image → Zoom → In [+]”, or put the mouse cursor in the region where you want to zoom in and use the keyboard shortcut (in Windows, use “Control” and +; in Mac OS, use “Command” and +).
6. To measure the parameters selected previously in step 2, go to ”Analyze → Measure” or use the keyboard shortcut (in Windows, use “Control plus M”, in Mac OS, use “Command plus M”. A new “Results” window will open displaying the measure values.
7. Close the image window and open a new image (select another image in case it’s already open).
8. Repeat steps 3 to 7 for each open image.
9. In the results window, select all the measurements (click on the Results window, then “Edit” → Select all”. Copy all measurements to the clipboard (“Edit” → Copy”).
10. Paste the results into a new Microsoft Excel file. Proceed with quantification and statistical analysis as desired.

## Expected outcomes

This protocol provides a reliable method (AID system) to efficiently degrade a protein of interest in *C. elegans*. Because *C. elegans* animals can be placed on NGM auxin (IAA or K-NAA) plates at different life stages, this system offers temporal control on protein depletion. Moreover, spatial control can be achieved by driving TIR1 expression using tissue-specific promoters (**Figure 1B**). Because AID is fused to a fluorescent reporter (e.g., *gfp, mNG*), depletion of the AID-tagged protein can be visualized *in vivo* with fluorescent microscopy, leveraging the optical transparency of *C. elegans*. The auxin treatment duration depends on the protein being depleted, but studies have demonstrated rapid depletion even after 30min of exposure to auxin ^3,13^. The degradation rate also depends on the concentration of auxin.

For the nuclear protein CFI-1 (ARID3), we observed efficient mNG::AID::3xFLAG::CFI-1 protein depletion in *C. elegans* neurons and muscle cells when L4 animals are placed on auxin plates for 2 days (**Figure 2A**). These animals were homozygous for the *ieSi57 [Peft-3::TIR1::mRuby::unc-54 3’ UTR, cb-unc-119(+)] II* line, which expresses TIR1 in all somatic cells(**Figure 2C**). Importantly, efficient CFI-1 depletion in neurons and muscle cells was observed upon either natural (IAA) or synthetic (K-NAA) auxin treatment (**Figure 2C**).

For the cytoplasmic protein Y47D3A.21 (DENR), we again observed efficient GFP::AID::3xFLAG::3xGAS:: Y47D3A.21 depletion upon either natural (IAA) or synthetic (K-NAA) auxin treatment (**Figure 2D**). These animals were homozygous for the pansomatic TIR1 line (*ieSi57).* Y47D3A.21 (*DENR)* depletion was observed not only by fluorescence microscopy, but also Western blotting (**Figure 2E**).

In our recent study^7^, we used the AID system for temporal and spatial depletion of CFI-1 in the nervous system. Specifically, we used a pan-neuronal promoter to drive TIR1 expression (*otTi28[unc-11prom8+ehs-1prom7+rgef-1prom2::TIR1::mTurquoise2::unc-54 3′UTR*) in all *C. elegans* neurons (**Figure 1B**). We performed a genetic cross between this strain and *cfi-1(kas16[mNG::AID::3xFLAG::cfi-1])*, resulting in homozygous *cfi-1(kas16[mNG::AID::exFLAGxcfi-1])*; *otTi28[unc-11prom8+ehs-1prom7+rgef-1prom2::TIR1::mTurquoise2::unc-54 3′UTR]* animals. Upon a 2 day-long exposure to natural auxin (IAA), we observed degradation of mNG::AID::3xFLAG::CFI-1 in motor neurons of the ventral nerve cord ^5^. Here, we repeated the natural auxin (IAA) treatment to compare it directly with synthetic auxin (K-NAA) treatment. As in our original study^7^, we witnessed significant mNG::AID::3xFLAG::CFI-1 depletion in nerve cord motor neurons – the levels of mNG fluorescence intensity are reduced upon either IAA or K-NAA treatment (**Figure 3A-C**). In these same animals, we also assessed mNG::AID::3xFLAG::CFI-1 depletion in head neurons and head muscle cells. Since TIR1 is provided only in neurons, we witnessed a significant reduction in the number of neurons expressing mNG::AID::3xFLAG::CFI-1, but no effect in head muscle cells, highlighting the tissue specificity of the otTi28 TIR1 line (**Figure 3C**). Importantly, similar effects were observed upon IAA or K-NAA treatment. Surprisingly, we did not detect robust mNG::AID::3xFLAG::CFI-1 depletion in neurons located at the tail (**Figure 3C**). By contrast, pansomatic TIR1 expression resulted in efficient mNG::AID::3xFLAG::CFI-1 depletion in tail neurons upon either IAA or K-NAA treatment (**Figure 2C**).

**Figure 3:**
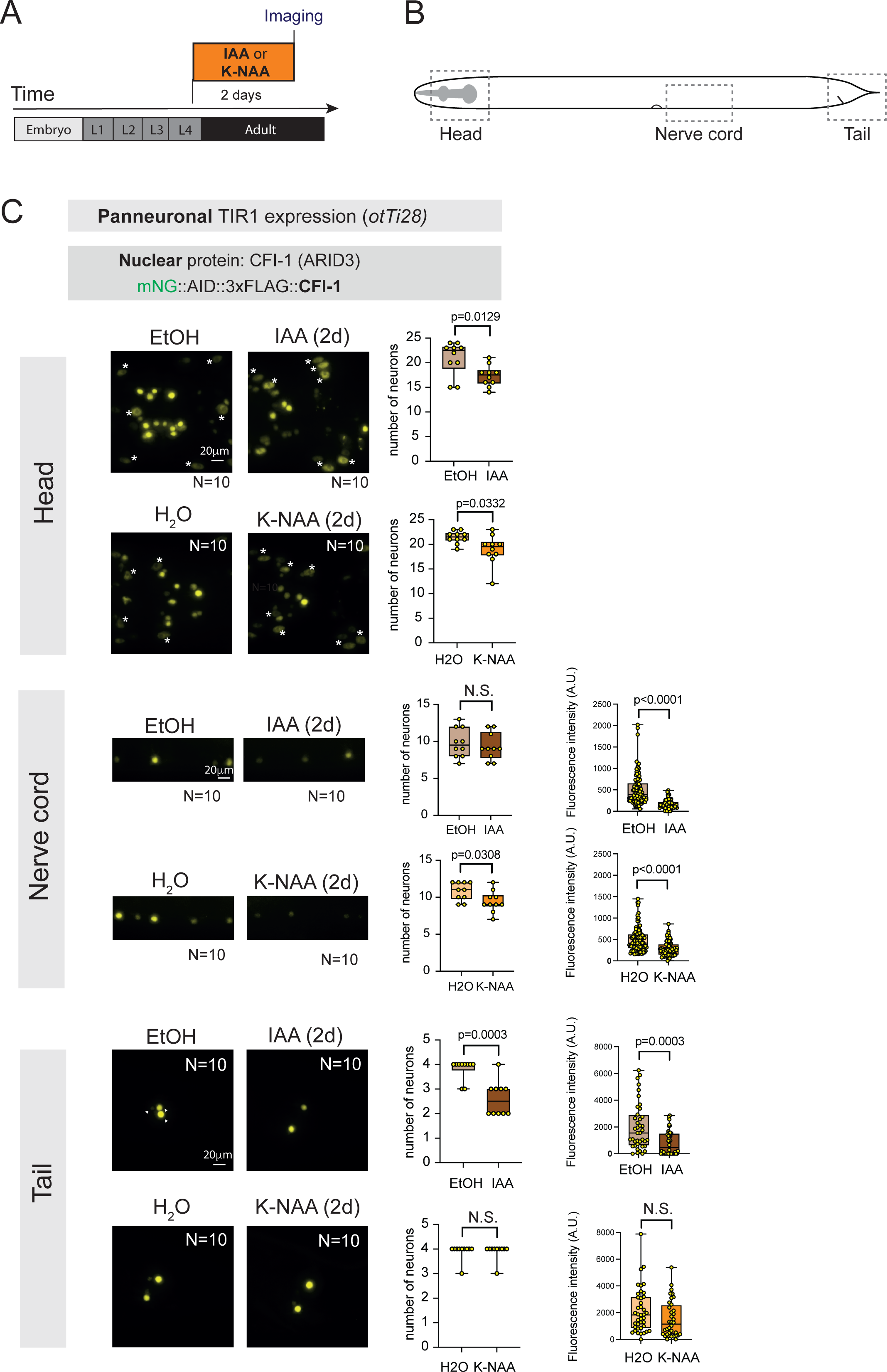
AID-mediated depletion of CFI-1 in the nervous system using natural (IAA) and synthetic (K-NAA) auxins. **A:** Schematic of auxin treatment. **B:** Schematic of *C. elegans* body. Boxed regions indicate images shown in panel C. **C**: Representative images showing mNG::AID::3xFLAG::CFI-1 expression in neurons and head muscle cells (asterisks) upon IAA or K-NAA treatment for two days. Controls: Two day-long exposure to either 100% ethanol or dH2O. Strain genotype: *cfi-1(kas16[mNG::AID::cfi-1]) I; otTi28 [unc-11prom8+ehs-1prom7+rgef-1prom2::TIR1::mTurquoise2::unc-54 3’UTR] X*. Images from the *C. elegans* head, ventral nerve cord, and tail are shown. Quantification of the number of neurons expressing mNG and fluorescence intensity are provided on the right. Student’s t test, N.S: Not significant.

More broadly, once efficient depletion of a protein is observed, this protocol is compatible with functional studies for virtually any *C. elegans* proteins. Three examples of applicability include:

1. **Molecular analysis of TF target genes** For temporally-controlled depletion of mNG::AID::3xFLAG::CFI-1, in our original study^7^, we placed L4 animals on NGM-auxin (4mM) and control plates for 2 days and examined expression of the cholinergic motor neuron-specific reporter *glr-4 (mScarlet)* with fluorescence microscopy at the adult (Day 2) adult stage. We observed that auxin-mediated CFI-1 depletion was accompanied by an increase in the number of motor neurons expressing *glr-4*. Hence, CFI-1 is required in the adult stage to repress *glr-4* in nerve cord motor neurons^7^.
2. This protocol can be combined with any *C. elegans* behavioral assays (e.g., harsh/gentle touch, osmotic avoidance, locomotion with automated worm tracking). Essentially, instead of preparing a microscope slide to evaluate gene expression, behavioral assays can be conducted, like for example the harsh touch used in the original study^7^. We note that K-NAA is dissolved in dH_2_O (or M9 buffer), whereas IAA is dissolved in ethanol. Because ethanol could confound certain behavioral or lifespan assays, K-NAA should be preferred for these types of experiments.
3. **Western blotting** In addition to microscopy, depletion of the protein of interest can be analyzed by a quantitative method, such as Western blotting (**Figure 2E**). To do so, place a large number of worms on auxin NGM plates (∼2 55mm plates). Upon lysis with SDS protein lysis buffer, proceed with determination of protein concentrations. Next, load samples (25-50 μg/ml) on an SDS gel, transfer to immunoblot and probe with protein-specific antibodies. One or two days after auxin (IAA) administration, we witnessed significant reduction of DENR protein levels by Western blotting (**Figure 2E**).

## Limitations

This protocol demonstrates an effective experimental set up for conditional degradation of AID-tagged proteins in *C. elegans.* Importantly, it also provides side-by-side comparisons of different auxins and TIR1-expressing lines that will benefit future studies of AID-mediated protein depletion in *C. elegans*. However, the AID system may not achieve complete protein elimination, and thereby the lack of a detectable phenotype has to be carefully interpreted. Success of this protocol depends: (a) on the generation of genetically modified *C. elegans* strains where the protein of interest is tagged with the AID degron, (b)TIR1 expression in specific cell types (i.e., neurons, muscles), and (c) exposure to natural (IAA) or synthetic (K-NAA) auxins. Further, several studies reported hypomorphic effects even in the absence of auxin^10,17^, likely due to: (a) tagging of the endogenous protein may interfere with its normal function, or (b) TIR1 may still be able to drive ubiquitination and subsequent degradation of the AID-tagged protein in the absence of auxin. Therefore, alternative approaches are recommended, such as tissue-specific RNAi, and conditional mutagenesis using the Cre/loxP recombination system^18,19^.

### Troubleshooting

#### Problem 1

Auxin induced protein depletion does not recapitulate expected outcomes or known phenotype. See Method details – step 2.

#### Potential solution

Auxin induced protein depletion can be affected by several factors such as level and timing of TIR1 expression and nature of protein of interest (stability).

- Before starting the auxin experiment, confirm the genotype of your strains.
- Include a positive control strain. If you do not observe the expected phenotype in the positive control, consider remaking the IAA working solution.

#### Problem 2

Worm growing on auxin plates does not look healthy. See Method details – step 2.

#### Potential solution

- Make sure to use NGM-auxin plates that are not older than 3 weeks from the date of preparation and stored in 4°C.
- Make sure you pick the worm from growing plate and in their healthier developmental stage. Picking worm from starved plate and transfer to auxin plate may illustrate unhealthy worm in the end of the experiment.
- You can also try the low range of auxin concentration (500 μM-1mM) and evaluate the best concentration works for your experiments.
- Before the experiment, let the OP50 on NGM plates dry completely.
- Natural auxin (IAA) slows the bacterial growth and can bacteria take longer to grow on IAA-NGM plates. K-NAA is suggested as it does not interfere with bacterial growth.

#### Problem 3

Auxin treatment is not effective for your protein of interest. See Method details – step 2.

#### Potential solution

- You can also try higher concentration of IAA or K-NAA solution (1mM-4mM). Depletion of AID-tagged protein of interest completely depends on the concentration of auxin. 1mM auxin could be low for some of proteins whose turnover is high, and the protein is highly stable.
- Upon high concentration of auxin (4 mM), it is recommended to use fresh and high concentrated OP50 every timing.
- UV-inactivated OP50 for high auxin concentration could be an alternative approach.
- As an alternative, you can spread/coat 50-100 ml of 4 mM auxin (in ethanol) on NGM plates using a spreader and let it complete dry in the dark.
- To avoid the use of high auxin concentration, we also recommend the AID2 system, which employs an OsTIR1(F74G) mutant and a synthetic auxin (5-Ph-IAA) ^10,11^. AID2 shows no detectable leaky degradation, requires a 670-times lower auxin concentration, and achieves even quicker degradation than the conventional AID system.
- You should carefully consider the duration of auxin exposure. You may want to try longer exposure times. However, the original study of AID in *C.elegans* demonstrated effective protein depletion in less than 4 hours. Therefore, when working with new strain, we recommend assessing a depletion efficiency at multiple time points.

#### Problem 4

AID::GFP knock-in in your transgene interfere with protein’s function. See Method details – step 2.

#### Potential solution

Endogenous tagging of the protein of interest with AID and a fluorescent protein (e.g., GFP) may interfere with protein function. If that is the case, tag the protein with a shorter epitope (e.g., FLAG) which may not disrupt protein’s function. Western blot can be used to measure the efficacy of protein knock-down by using anti-FLAG antibody.

#### Problem 5

Worms are still mobile in the anesthetizing solution. The cause must be an insufficient concentration of the anesthetizing solution. See Method details – step 5a.

#### Potential solution

Increase the concentration of NaN_3_, however avoid a highly concentrated solution as it may kill the worms.

#### Problem 6

High background. The cause may be an excess of bacteria transferred when picking the worms. See Method details – step 5b.

#### Potential solution

Minimize transfer of bacteria while picking the worms.

#### Problem 7

Worms burst frequently. This may happen for several reasons: See Method details – step 5c.

- The samples waited too long for imaging after preparation.
- The volume of the anesthetizing solution is too low, or its concentration is too high.
- The concentration of the agarose solution used for the pad is too high.

#### Potential solution

The pads help worms stay moist and avoid desiccation, however samples should be imaged within 30 minutes. If worms burst within this timing, the problem must be either the agarose pad or the anesthetizing solution. Lower the concentration of the agarose or the anesthetizing solution. Try increasing the volume of anesthetizing droplets.

#### Problem 8

Worms scatter too much on the agarose pad. If the agarose pad is too large and the volume of anesthetizing solution is too high. See Method details – step 5a.

#### Potential solution

Drop a smaller volume of agarose solution to make the pad. Place anesthetizing solution onto the coverslip.

#### Problem 9

Cannot focus the sample. The cause may be a dirty objective lens. See Method details – step 10 and 11.

#### Potential solution

Make sure the lenses are clean. Clean only with lens cleaner. Use immersion oil for higher magnification objectives (usually 40x and above).

#### Problem 10

How to distinguish fluorescence from autofluorescence.

#### Potential solution

Since autofluorescence is common in *C. elegans,* it may interfere with detection of fluorescently tagged proteins fused to AID, such as mNG::AID::3xFLAG::CFI-1. However, autofluorescence in *C. elegance* mostly comes from granules in the intestine, and is more evident in the green channel (488 nm excitation). It becomes more evident in adult animals. Hence, imaging younger animals may help.

## Resource availability

### Lead contact

Further information and requests for resources and reagents should be directed to and will be fulfilled by the lead contact, Paschalis Kratsios.

### Materials availability

The protocol described here did not generate any new method or unique reagents. All *C. elegans* strains used in this protocol are available from the lead contact and/or CGC (https://cgc.umn.edu).

### Data and code availability

- All data generated for or analyzed in this study are contained in the manuscript and supporting files. Microscopy data will be shared by the lead contact upon request.
- Any additional information required to analyze or reanalyze data in this manuscript is also available upon request.
- No code was generated to analyze the data.

## Acknowledgments

We thank the *Caenorhabditis* Genetics Center (CGC), which is funded by NIH Office of Research Infrastructure Programs (P40 OD010440), for providing the CA1200 strain, as well as the lab of Dr. Oliver Hobert for the OH14930 strain. This work was funded by a Pilot Grant

Award from the Safadi Program of Excellence in Clinical and Translational Neuroscience to N.S., a Swiss National Science Foundation (SNSF) postdoctoral fellowship to F.M., and two NIH grants (R21 NS108505, R01 NS118078) to P.K.

## Author contributions

Conceptualization and Methodology: N.S., F.M., P.K., Investigation, N.S., F.M.; Writing – original draft, review and editing, N.S., F.M., P.K., Funding acquisition, Resources, and supervision, P.K.

## Figure legends

**Supplementary Figure 1:**
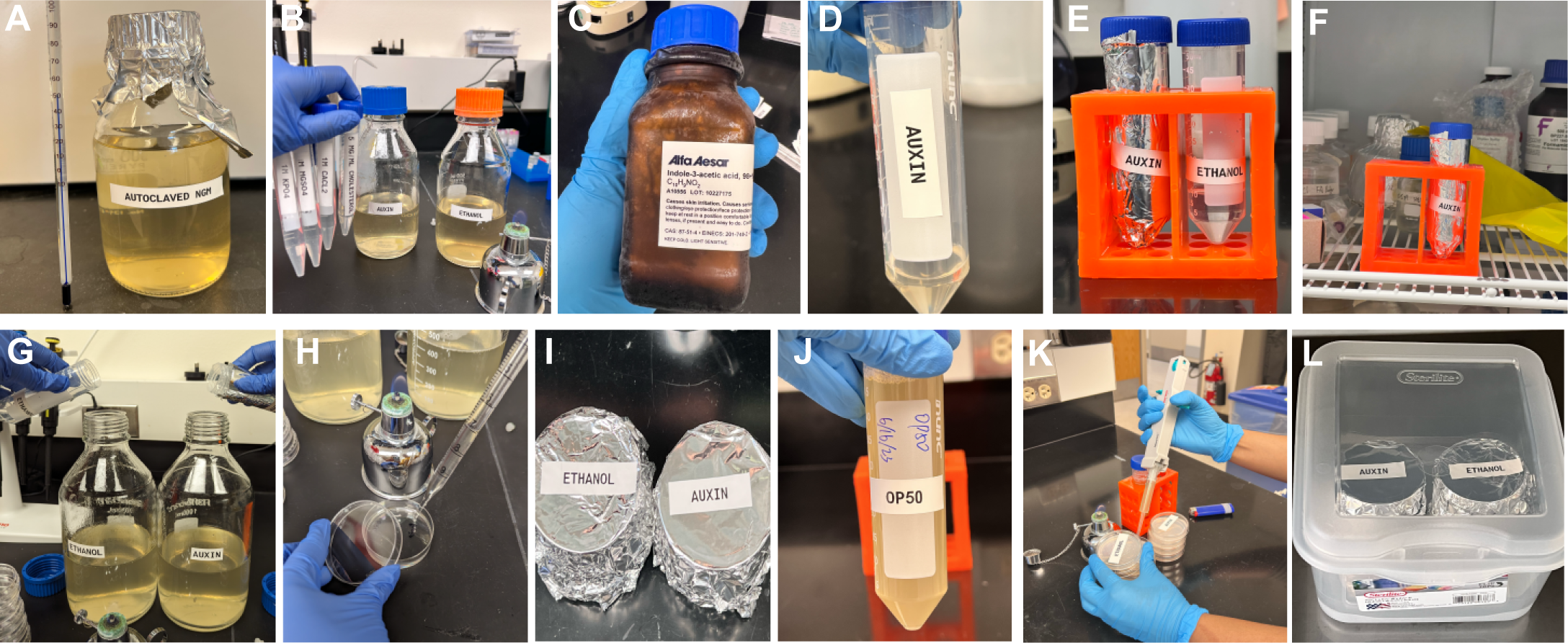
Preparation of NGM and auxin/ethanol coated NGM plates. A. Prepare nematode growth medium (NGM). Autoclave the NGM and let it cool down at 55 °C (water bath or hot plate) until ready to use. B. Add buffers and cholesterol to the autoclave NGM and divide into two bottles for ethanol and auxin. C. Prepare fresh auxin (stored at -20 °C). D. Prepare the 400 mM auxin (IAA or K-NAA) stock dissolved in ethanol (for IAA) or dH_2_O (for K-NAA). The final solution should be crystal clear as shown above in E. E. Shield the auxin with aluminum foil throughout. F. 400 mM auxin stock solution can be stored at 4 °C for up to a week. G. Add the 400 mM auxin to the cooled down (55 °C) NGM to make the final concentration of 4 mM auxin as well as an equal volume of ethanol (or dH_2_O) to the NGM as a control. H. Pour 10 mL of auxin-NGM and ethanol-NGM into the respective 55 mm culture petri dish. I. Shield the plates of ethanol and auxin and let them dry for a day. J. Use concentrated OP50 to allow good bacterial growth on auxin and ethanol plate. K. Next day seed the plate with 200 μL of OP50 and let them dry for 2 more days at the room temp and in a sterile place. L. Shield the ethanol and auxin plates, that can be stored at 4 °C for up to a month.

**Supplementary Figure 2:**
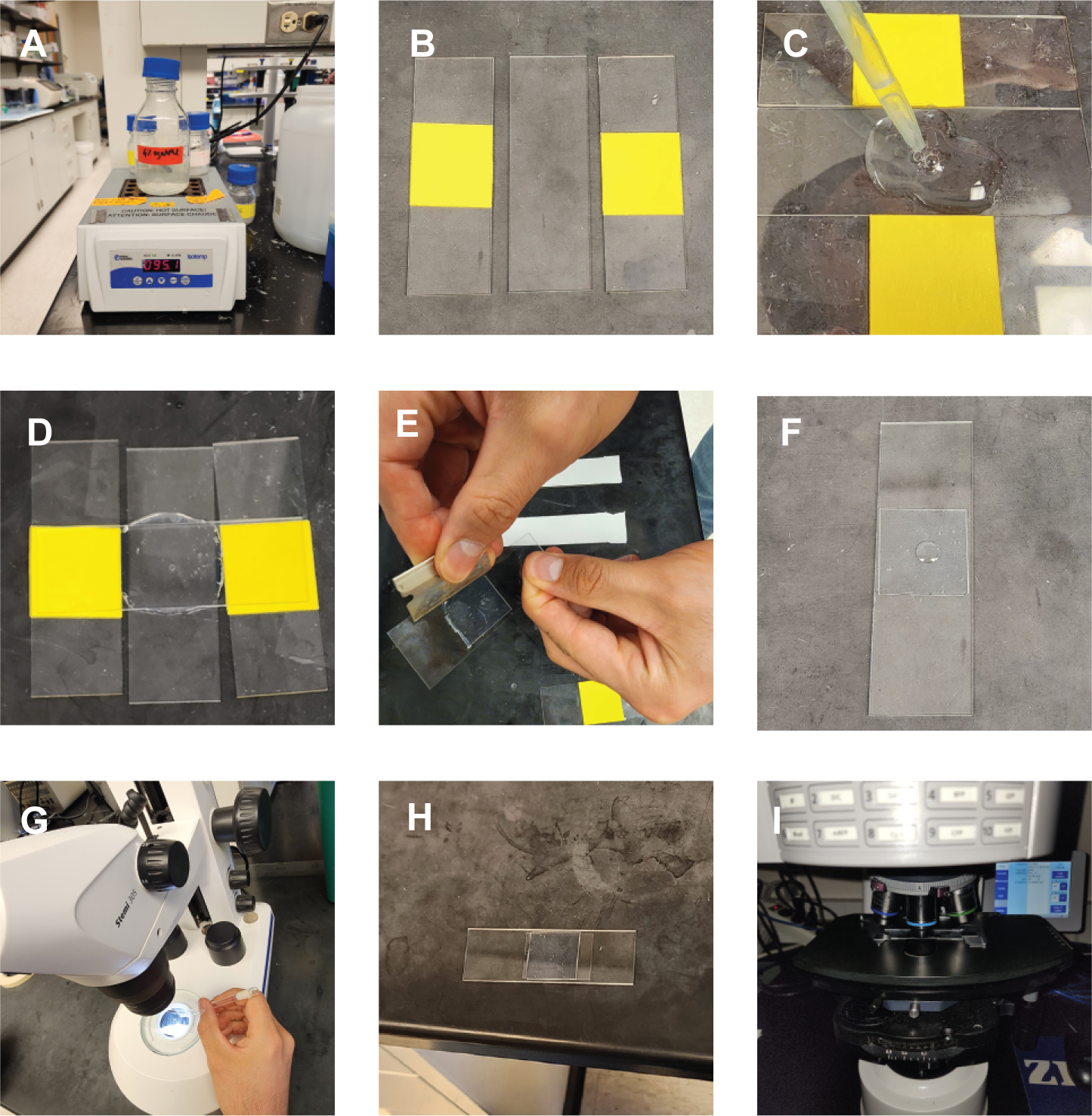
Preparation of imaging slides. (A) Prepare 4% agarose solution. Keep the melted agarose on a heat block, at approximately 95 °C. (B) Place the support slides on each side of a blank slide. (C) Place one drop (∼600L) of warm 4% agarose onto the blank slide using a laboratory pipette with a cut blue tip. (D) Quickly place a second blank slide on the top of the agarose drop, at a 90° angle to the three slides, and gently press the edges of this slide such that the agarose spreads uniformly. Let the agarose dry for a minute. (E) Separate the glass slides without perturbing the agarose pad. Use a razor blade to trim excessive parts and adjust the agarose pad. (F) Place a 10 μL droplet of 100mM sodium azide (NaN3) on the agarose pad. (G) Pick the worms to be imaged under the dissection scope and put them in the NaN3 droplet. Gently cover the droplet with a coverslip. (I) Place imaging slide on the microscope stage with the glass coverslip facing the objective lens.

**Supplementary Figure 3:**
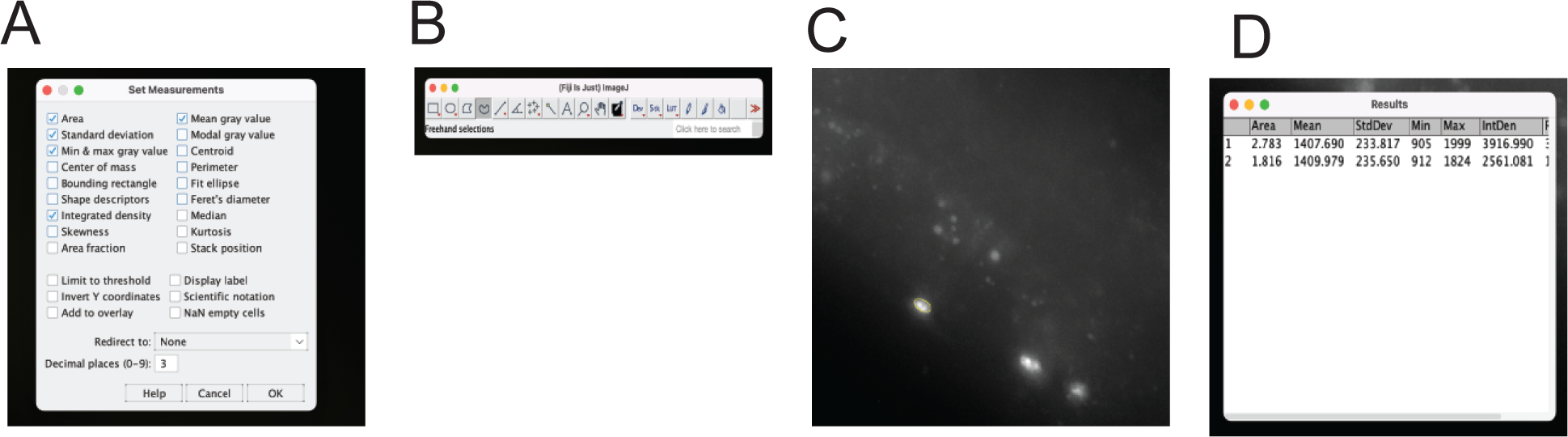
Image analysis with ImageJ. (A) Check the desired parameters and click OK to close the window. (B) In the ImageJ window, click on the freehand selection tool. (C) Carefully outline the region to be quantified and execute the measure function. (D) A new “Results” window will open displaying the measures values.

